# Membrane Stretch Gates NMDA Receptors

**DOI:** 10.1101/2022.01.17.476675

**Authors:** Sophie Belin, Bruce A. Maki, James Catlin, Benjamin A. Rein, Gabriela K. Popescu

## Abstract

N-Methyl-D-aspartic (NMDA) receptors are excitatory glutamate-gated ion channels. Their activation is essential for the normal development, maintenance, and plasticity of excitatory synapses in the central nervous system. They function as glutamate-gated Ca^2+^-permeable channels, require glycine as co-agonist, and can be modulated by myriad of diffusible ligands and cellular cues, including mechanical stimuli. Previously, we found that in cultured astrocytes, shear stress initiates NMDA receptor-mediated Ca^2+^ entry in the absence of added agonists, suggesting that in addition to being mechanosensitive, NMDA receptors may be mechanically activated. Here, we used controlled expression of recombinant receptors and non-invasive on-cell single-channel current recordings to show that gentle membrane stretch can substitute for the neurotransmitter glutamate in gating NMDA receptor currents. Notably, stretch-activated currents preserved the hallmark features of the glutamate-gated currents, including glycine-requirement, large unitary conductance, high Ca^2+^ permeability, and voltage-dependent Mg^2+^ blockade. Further, we found that the stretch-gated current required the receptor’s intracellular domain, which may suggest a force-from-filament sensing mechanism. These results are consistent with the hypothesis that mechanical forces can gate NMDA receptor currents even in the absence of synaptic glutamate release, which has important implications for understanding mechanotransduction and the effect of mechanical forces on cells of the central nervous system.

**Highlights:**

- Membrane stretch gates NMDA receptor currents in the absence of the neurotransmitter glutamate.
- Stretch-gated currents maintain the characteristic features of glutamate-gated currents, including glycine requirement, Ca^2+^ permeability, and voltage-dependent Mg^2+^ block.
- Gating of NMDA receptor by membrane stretch requires the receptor’s intracellular domain.
- Mild stretch of neuronal membranes gate native NMDA receptor currents.

**Summary:** Membrane stretch gates NMDA receptor currents in the absence of neurotransmitter. Stretch-gated currents have the biophysical hallmarks of the glutamate-gated currents including requirement for glycine, large Na^+^ conductance, high Ca^2+^ permeability, and voltage-dependent Mg^2+^ block.

## Introduction

Cells of the central nervous system (CNS) experience endogenous and environmental mechanical forces *in vivo*, and respond to osmotic and atmospheric pressure *ex vivo* (Bliznyuk, Hollmann, & Grossman, 2020; Koser et al., 2016; Tyler, 2012). Mechanical stimuli affect several neurophysiological processes including neuronal firing, vesicle fusion, dendritic spine formation, and synaptic activity (Hill, 1950; G. H. Kim, Kosterin, Obaid, & Salzberg, 2007; Korkotian & Segal, 2001; Star, Kwiatkowski, & Murthy, 2002; Ucar et al., 2021). However, the mechanism of mechanotransduction in the CNS remains poorly understood largely due to experimental, technological, and theoretical challenges unique to examining the effect of mechanical forces in biological tissues. Among these obstacles are the omnipresence of mechanical cues, their diverse three dimensional and dynamic actions, the variety of macromolecules that participate in mechanotransduction, and the multiplicity of mechanisms by which transducers sense and respond to mechanical stimuli (Cox, Bavi, & Martinac, 2019; Kefauver, Ward, & Patapoutian, 2020; Le Roux, Quiroga, Walani, Arroyo, & Roca-Cusachs, 2019).

On a millisecond time-scale, mechanotransduction in the nervous system is mediated by mechanically-activated and mechanically-sensitive ion-channels (Cox et al., 2019; Kefauver et al., 2020). Mechanically-activated channels are membrane proteins dedicated to scanning the environment for mechanically-encoded information; they represent the molecular basis for a wide array of mechanosensory processes including hearing, touch, and proprioception; and are critical for normal development and adaptation throughout life (Murthy, Dubin, & Patapoutian, 2017; Walsh, Bautista, & Lumpkin, 2015). On the other hand, a large swath of ion channels whose primary physiological function is to respond to electrical and chemical signals, while not directly gated by mechanical stimuli, are mechanosensitive. These channels mediate much of the CNS mechanotransduction and are essential to how mechanical forces influence the normal development and functioning of the brain and spinal cord, and also how they initiate or aggravate acute and chronical neuropathologies (Tyler, 2012).

N-methyl-D-aspartate (NMDA) receptors are glutamate-gated channels with demonstrated mechanosensitivity (Johnson, Battle, & Martinac, 2019). NMDA receptors mediate excitatory transmission and plasticity in CNS and are critical for the normal physiology of excitatory synapses; moreover, their overactivation mediates glutamate excitotoxicity, which has been implicated as a causal factor in several neuropathologies. Ambient pressure, membrane stretch, and membrane lipid composition modulate their agonist-gated currents in native preparations, in heterologous systems, and in artificial lipid bilayers (Casado & Ascher, 1998; Fagni, Zinebi, & Hugon, 1987; Kloda, Lua, Hall, Adams, & Martinac, 2007; Miller, Sarantis, Traynelis, & Attwell, 1992; Nishikawa, Kimura, & Akaike, 1994; Paoletti & Ascher, 1994). In addition to mechanosensitivity, we reported recently that shear stress, as applied by shear microfluidic flow onto cultured astrocytes, elicits NMDA receptor-mediated Ca^2+^ influx in the absence of glutamate, suggesting that mechanical stimuli *per se* can gate NMDA receptor currents (Maneshi et al., 2017). This observation has important implications for a possible role of NMDA receptors in mechanotransduction during the normal development and function of the CNS (Goriely et al., 2015; Heuer & Toro, 2019; Tyler, 2012); and also in severe neuropsychiatric pathologies, including those associated with acute traumatic brain and spinal cord injuries, chronic traumatic encephalopathy, shaken baby syndrome, and episodic edema or tumor growth (Bonnier, Mesples, & Gressens, 2004; Shively, Scher, Perl, & Diaz-Arrastia, 2012; Sloley et al., 2021). Therefore, we undertook the work reported here to investigate this novel observation in more depth.

NMDA receptors are widely expressed in neuronal and glial cells of the CNS as heterotetramers of two obligatory GluN1 subunits and two additional subunits, from the related families of GluN2 (A – D) and GluN3 (A, B) subunits. Of the glutamate-binding GluN2 subunits, GluN2A predominates in adult animals and at mature synapses, whereas GluN2B is expressed mostly in juvenile animals and at immature synapses (Goebel & Poosch, 1999; Monyer et al., 1992; Paoletti, Bellone, & Zhou, 2013). Within the larger family of ionotropic glutamate receptors, NMDA receptors are unique because in addition to glutamate, they require glycine as an obligatory gating co-agonist. They also have characteristically large unitary conductance, high Ca^2+^ permeability, and voltage-dependent Mg^2+^ block (Hansen et al., 2018). These distinctive biophysical properties of the glutamate-gated current are essential for many of the critical functions of NMDA receptors in health and disease (Hansen et al., 2018; Iacobucci & Popescu, 2017).

Based on our previous observation that shear force can elicit native NMDA receptor-dependent Ca^2+^ fluxes in primary culture of astrocytes in the absence of agonist (Maneshi et al., 2017), here we investigate mechanically-activated NMDA receptor currents using a recombinant system and single-channel patch-clamp current recordings. We found that, as with sheer stress in astrocyte, gentle suction applied to the membrane patch elicited currents from recombinant NMDA receptors expressed in HEK cells in the absence of the neurotransmitter glutamate. Importantly, the stretch-gated current maintained the characteristic biophysical properties of the glutamate-gated current, including requirement for glycine, high unitary conductance, Ca^2+^-permeability, and voltage-dependent Mg^2+^ blockade. In addition, we found that the C-terminus of NMDA receptors is required for gating by membrane stretch, which may suggest a force-from-filament mechanism of force transmission.

## Materials and Methods

### Cells and receptor expression

HEK293 cells (American Type Culture Collection number CRL-1573) were grown and maintained in Dulbecco’s modified Eagle’s medium along with 10% fetal bovine serum (FBS, Gibco) and 1% penicillin/streptomycin. Cells were grown to 80% confluency, and passages 24 - 31 were used for transfections. Cells were transfected transiently via the Ca^2+^-phosphate method using pcDNA3.1 (+) plasmids encoding for rat GluN1-1a (P35439-1), GluN2A (Q00959) and GFP (P42212) in a 1:1:1 ratio. When indicated, plasmids encoding for GluN1-1a and GluN2A were replaced by plasmids encoding CTD-truncated GluN1-a (GluN1-a 838stop) and CTD-truncated GluN2A (GluN2A 844stop), provided by Dr. Westbrook (Krupp, Vissel, Thomas, Heinemann, & Westbrook, 1999b, 2002). Alternatively, when indicated, the GluN2A-encoding plasmid was substituted with plasmids expressing GluN2B (Q00960), GluN2C (Q00961), or GluN2D (Q62645). Cells were incubated with the DNA mixture for 2 hours, were washed twice with phosphate buffer saline (PBS), and incubated in growth medium supplemented with 2 mM MgCl_2_, to prevent excitotoxicity. They were used for electrophysiological recordings within 24 hours.

### Culture of Dissociated Hippocampal Neurons

Low-density cultures of acutely dissociated hippocampal neurons were prepared from Sprague-Dawley rat embryos (Envigo) at embryonic day 18 (E18) with minor adjustments from previously described methods (Misonou & Trimmer, 2005). Briefly, a pregnant rat was euthanized in a CO_2_ chamber and decapitated, and the uterus was surgically removed. Embryos were decapitated, and the hippocampi were removed and placed in ice-cold dissecting solution containing HBSS supplemented with 4 mM sodium bicarbonate (Sigma), 10 mM HEPES (Sigma) and 1% penicillin/streptomycin (Corning). Cells were enzymatically dissociated with 0.25% trypsin (20 min at 37°C), and then gently triturated with a stereological pipette and filtered through a 40 mm strainer (BD Falcon, Franklin Lakes, NJ). Dissociated cells were counted and plated at a density of 100,000 cell/cm² onto glass cover-slips precoated with poly-D-lysine (Corning) in plating media containing MEM (Gibco) supplemented with 10% FBS, 0.6 % glucose (Sigma), 2 mM GlutaMAX (Gibco), 1 mM sodium pyruvate (Sigma), and 1% penicillin/streptomycin. Cells were allowed to adhere to plates for several hours after which the medium was gently replaced with Neurobasal A medium (Gibco) supplemented with B27 (Gibco) and 2 mM GlutaMAX. At 3 days in vitro (DIV), the proliferation of non-neuronal cells was inhibited by adding 5µM arabinofuranosylcytosine (Sigma). Neurons were used for electrophysiological experiments between 7 and 30 DIV.

### Electrophysiology

To maintain consistency in seal formation with minimal mechanical disruption to the patch, we used the following procedure. Prior to entering the bath, we applied slight positive pressure (5 mmHg) through the recording pipette with a high speed pressure-clamp system (HSPC-1, ALA Scientific, Farmingdale, NY) (McBride & Hamill, 1993, 1999). Electrical resistance through the pipette (20 ± 5 MΩ) was monitored by observing the amplitude of the current elicited by a test voltage-pulse. After contacting the cell, the positive pressure was released to 0 mmHg, and slight suction (−5 mmHg) was applied to initiate gentle seal formation onto the cellular membrane, which was monitored as an increase in pipette resistance. Finally, after obtaining a high-resistance seal, we released the negative pressure and applied +100 mV to the patch to visualize the activity of NMDA receptors at 0 mmHg, as inward currents.

To examine the dependency of channel activity on the level of applied pressure, cells were bathed in PBS; after seal formation, we applied pressure in increments of 10-mmHg, and recorded activity for periods lasting ∼5 minutes for each pressure level, over the indicated range. When specified, a 5-minute recovery step was recoded after relaxing the pressure to 0 mmHg. Channel activity was evaluated in cell-attached patches obtained with pipettes filled with (in mM) 150 NaCl, 2.5 KCl, 10 HEPBS, 1 EDTA, pH 8.0 (NaOH) and the indicated agonists glutamate (1 mM), glycine (0.1 mM), or NMDA (0.1 mM), as previously described (Hamill, Marty, Neher, Sakmann, & Sigworth, 1981; Maki, Cummings, Paganelli, Murthy, & Popescu, 2014). Solutions lacking glycine were prepared using double-distilled deionized ultrapure water (Fisher Scientific, Hampton, NH) to prevent contamination with glycine (Cummings & Popescu, 2015).

To examine the effect of pressure on the receptor’s conductance, Ca^2+^ permeability, and voltage dependency of its Mg^2+^ blockade, cells were bathed in a high K^+^-based bath solution (in mM): 142 KCl, 5 NaCl, 1.8 CaCl2, 1.7 MgCl2, 10 HEPBS, pH 7.2 (with KOH) to collapse the physiological membrane potential of HEL cells, which is ∼10 mV (Borschel et al., 2012). Pipette solution was: 150 NaCl, 2.5 KCl, 10 HEPBS, 1 EDTA, 10 tricine, pH 8.0, and glycine (0.1 mm) and glutamate (1 mM) as indicated. Ca^2+^ and Mg^2+^ were added as chloride salts and were buffered to the indicated free concentration according MAXC software (maxchelator.standford.edu). After seal formation, we applied sustained suction (−40 mmHg) and varied the applied voltage in 20-mV increments, each lasting 1 minute, over the +100 mV to +20 mV range.

All current traces were filtered (10 kHz), amplified (Axopatch 200b) and then sampled (40 kHz) and stored as digital files using QuB software (www.qub.buffalo.edu).

### Data analysis

Current traces were inspected visually off line and only low-noise, stable-baseline recordings were selected for analyses. Traces were initially processed to correct for spurious noise events or minor baseline drifts (Maki et al., 2014). Corrected traces were idealized separately for each applied pressure within the QuB suite for kinetic analyses, with SKM algorithm after applying a digital filter (12 kHz) (Qin, 2004). We estimated the open probability (nPo) in each trace according to the following relationship:

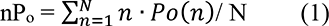

Where Po is the open probability of each channel, n is the indeterminate number of channels in each patch, and N the minimum number of channels in each patch, estimated as the maximum number of overlapping unitary currents (simultaneous openings) observed in the condition producing maximal activity. Values for nP_o_ were obtained by averaging activity in each 5-minute segment, and were considered non-zero for a threshold of >1,000 events.

Unitary channel conductance (γ) and reversal potentials (E_rev_) were estimated from linear fits to the unitary current-voltage relationship measured over a one-minute period. Ca^2+^ permeability was estimated as a function of the measured Ca^2+^-induced shifts in E_rev_ using the Lewis Equation (Lewis, 1979).

### Statistics

Results are illustrated as mean ± SEM with a minimum of three replicates per condition. Statistical analyses were performed using one-way ANOVA with Bonferroni’s multiple comparison test or Student’s *t*-test relative to control as indicated.

## Results

### Membrane stretch substitutes for glutamate in gating NMDA receptors

To examine whether NMDA receptors are simply mechanically sensitive or whether they can be gated by mechanical forces in the absence of neurotransmission, we expressed recombinant GluN1/GluN2A receptors in HEK293 cells and recorded inward Na^+^ currents from cell-attached patches, while gently varying the pressure applied through the recording pipette in 10 mmHg-increments over the −40 mmHg to +40 mmHg range. These pressures are typical for the activation of physiologic mechanotransducers such as Piezo channels (Coste et al., 2012; S. E. Kim, Coste, Chadha, Cook, & Patapoutian, 2012). Observing NMDA receptor activity over long periods is necessary to reduce patch-to-patch variability due to modal gating, which for NMDA receptors occurs on a minutes time scale (Borschel et al., 2012; G. Popescu & Auerbach, 2003). Therefore, at each pressure level, we recorded activity continuously for 5 minutes.

When the recording pipette included supra-saturating levels of the neurotransmitter glutamate (1 mM; Kd, 3 μM) (G. Popescu, Robert, Howe, & Auerbach, 2004) and the obligatory co-agonist glycine (0.1 mM; Kd, 2.5 μM) (Cummings & Popescu, 2015), applying +100 mV through the pipette produced large inward unitary currents (8 – 10 pA) indicative of channel activation, at all levels of applied pressure tested (Figure 1A, top traces Glu/Gly). Often, overlapping openings were apparent, indicating that multiple active channels were trapped in the recorded patch. In these conditions, neither negative nor positive pressure altered channel activity. When glycine was omitted, we observed only minimal and sporadic currents (<1,000 events per 5-min segment), regardless of whether glutamate was present or not, and applying either negative or positive pressure did not alter this low baseline-activity (Figure 1A, middle traces Glu/-). However, when glycine was present, negative but not positive pressure gated substantial current in the absence of glutamate (Figure 1A, bottom traces -/Gly). The suction-gated current increased with increasing pressure in a consistent manner, although the magnitude of the effect varied. On average, suction (−40 mmHg) increased GluN1/GluN2A channel activity (nPo) from 0.10 ± 0.05 to 0.56 ± 0.18 (n = 6, P < 0.03) (Figure 1A, B). This result demonstrates that suction alone can gate GluN1/GluN2A receptors, and therefore that it is possible to open NMDA receptors mechanically, in the absence of neurotransmission.

**Figure 1.**
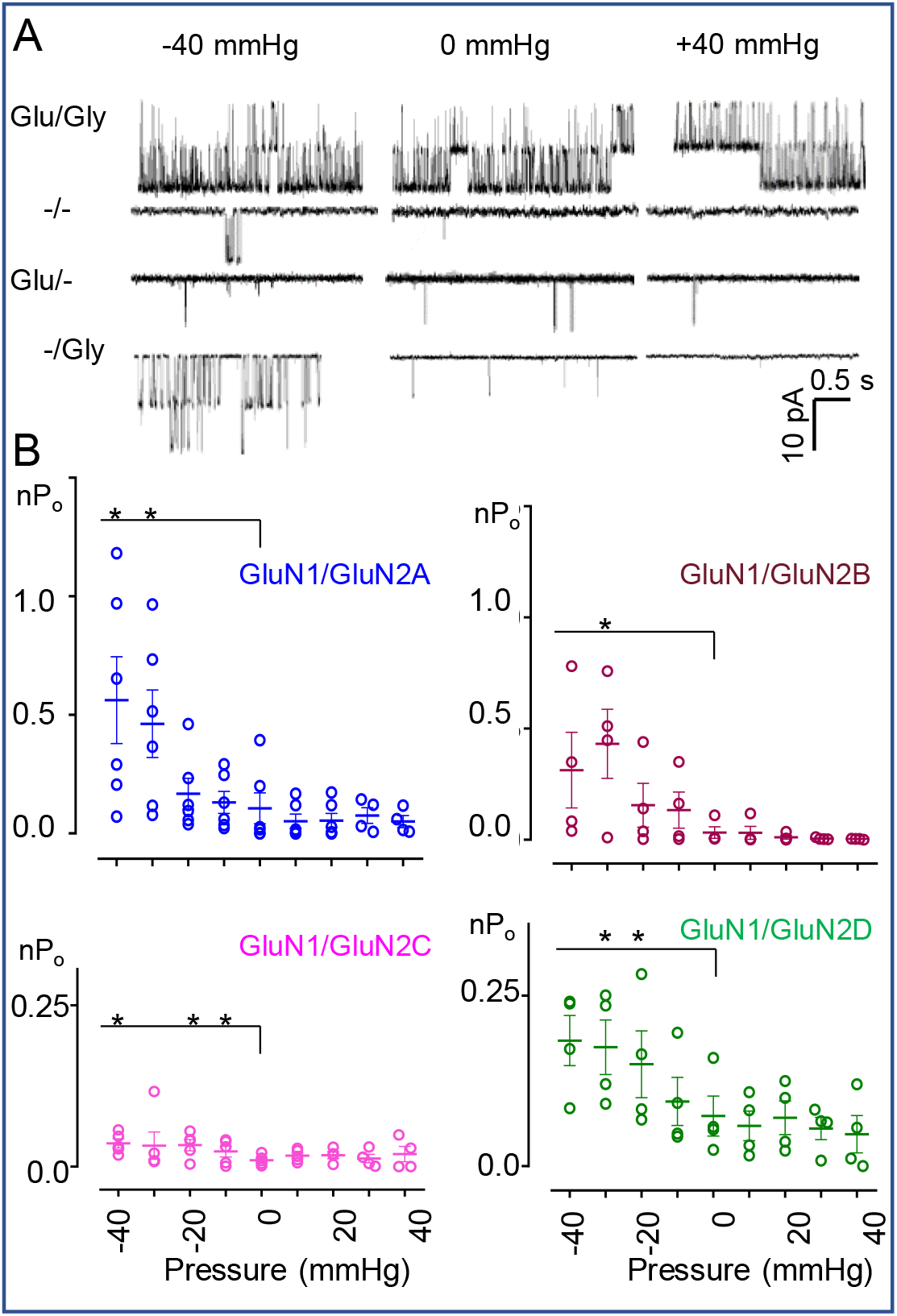
Effect of gentle hydrostatic pressure on currents recorded from recombinant NMDA receptors. **(A)** Current traces recorded from cell-attached patches expressing recombinant GluN1/GluN2A receptors with +100 mV applied through the recording pipette. Downward deflections illustrate inward Na^+^ currents at the indicated pressure levels, in the presence (+) or absence (−) of glutamate (Glu, 1 mM) and/or glycine (Gly, 0.1 mM). **(B)** Summary of response-dependency on applied pressure for each GluN2 subtype in the presence of glycine (0.1 mM) with no glutamate added. *, P < 0.05 (paired Student’s t-test).

To ascertain whether pressure can gate currents from other members of the NMDA receptor family, we co-expressed GluN1 with GluN2B, GluN2C or GluN2D subunits in HEK293 cells and recorded single-channel inward Na^+^ currents from cell-attached patches with pipettes containing glycine (0.1 mM) but not glutamate. As with the adult GluN2A isoform, we observed a selective increase in channel activity with negative pressure, and no effect with positive pressure of similar magnitude (Figure 1B). On average, application of −40 mmHg increased channel activity (nPo) as follows: for GluN2B from 0.03 ± 0.02 to 0.31 ± 0.17 (P > 0.05, n = 4); for GluN2C, from 0.010 ± 0.003 to 0.036 ± 0.007 (n = 4, P < 0.02); and for GluN2D, from 0.07 ± 0.03 to 0.18 ± 0.03 (P > 0.05, n = 4). Although suction increased the channel open probability in all patches, the average increase was not consistently statistically significant (Figure 1B). This may be explained in part by the documented intrinsically higher kinetic variability of GluN2B- and GluN2C-containg subunits, (Amico-Ruvio & Popescu, 2010; Khatri et al., 2014), and the low open probability of GluN2D-containing receptors, which makes detection more challenging (Vance, Hansen, & Traynelis, 2013). Overall, these results support the hypothesis that mechanical forces in addition to modulating the glutamate-gated current, can by themselves provide the energy necessary to shift the receptor’s closed-to-open equilibrium and produce a substantial increase in open probability. We focused next on GluN1/GluN2A receptors, which generally present more robust and reliable responses (Borschel et al., 2012).

Mindful of the many sources that can contribute to the variability of the observed effect, we aimed to reduce possible effects due to cellular changes over the long recording period necessary to cover the 80 mmHg-range investigated with the protocol above. For this, we shortened the observation period by limiting measurements to the effects of negative pressure, and added a 5-minute recovery step, to verify the reversibility of the pressure-dependent effect. As in the first set of experiments, with this shorter protocol we found that negative pressure had no effect on channel activity in the absence of glycine, or in the presence of saturating concentrations of glycine and glutamate (Figure 2, Table 1). However, when glutamate was omitted, −30 mmHg increased the observed current (nPo) from 0.015 ± 0.007 to 0.061 ± 0.026 (n = 6, P = 0.02); and −40 mmHg further increased nPo to 0.076 ± 0.022 (n = 4, P = 0.03). On average, the current gated by −40 mmHg represented ∼15% of the maximal glutamate-gated current in the same conditions (0.44 ± 0.14, n = 4). This level of activity is on par with that reported for extrasynaptic receptors activated by synaptic glutamate spill-over or by glutamate leak from injured neurons (Moldavski, Behr, Bading, & Bengtson, 2020). This level of current may be physiologically significant if the stretch-gated currents maintain the biophysical properties of glutamate-gated currents, especially their large unitary conductance, high Ca^2+^ permeability, and voltage-dependent Mg^2+^-block. Therefore, we next examined these critical biophysical properties of the stretch-gated current.

**Figure 2.**
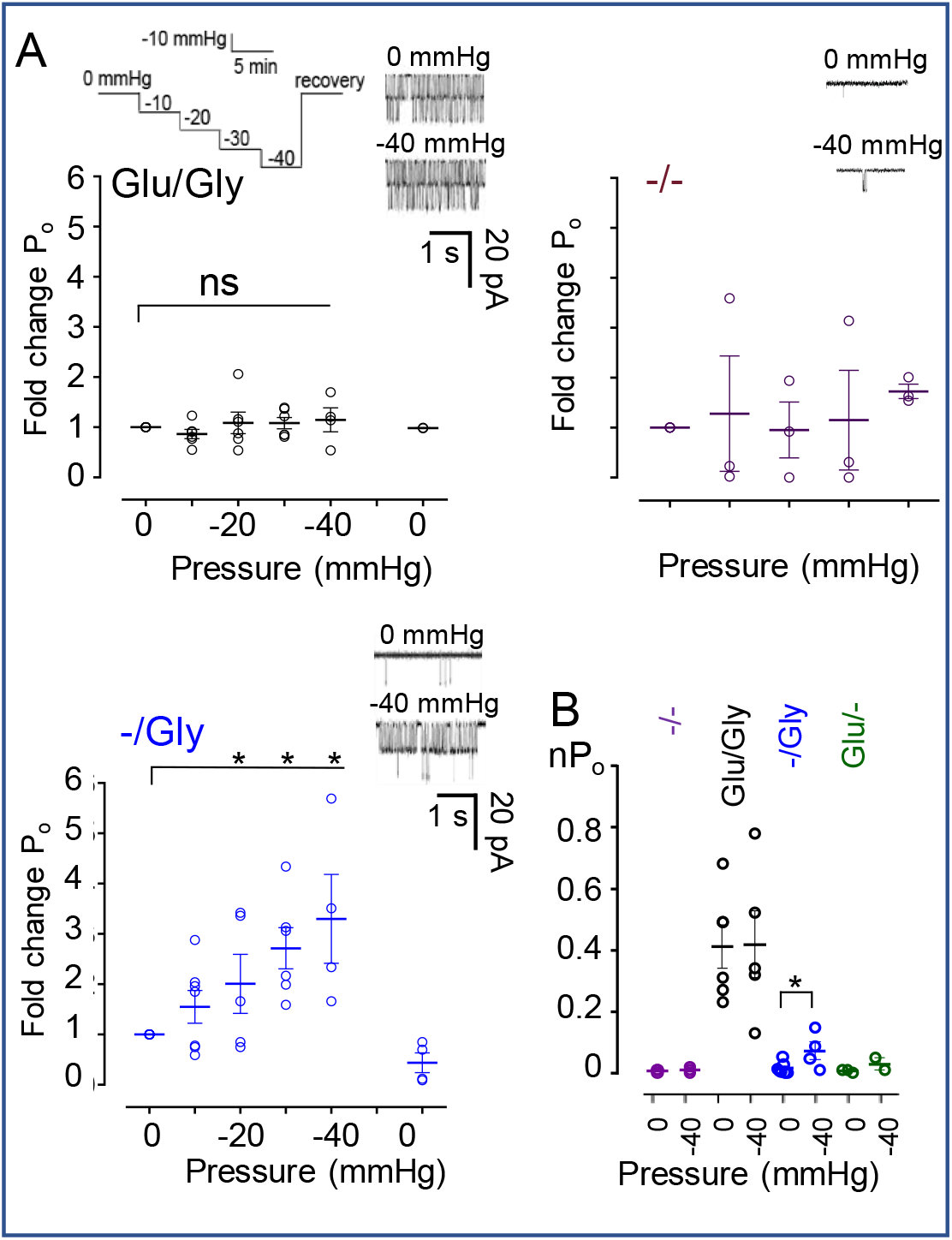
Gentle suction agonizes currents from NMDA receptors. **(A)** Effect of suction, applied with the illustrated protocol, on GluN1/GluN2A currents recorded with +100 mV in the cell-attached pipette, with the indicated diffusible agonists (Glu 1 mM, Gly 0.1 mM). **(B)** Summary of the effects of suction on receptor activity. *, P < 0.05 (Student’s t-test).

**Table 1.**
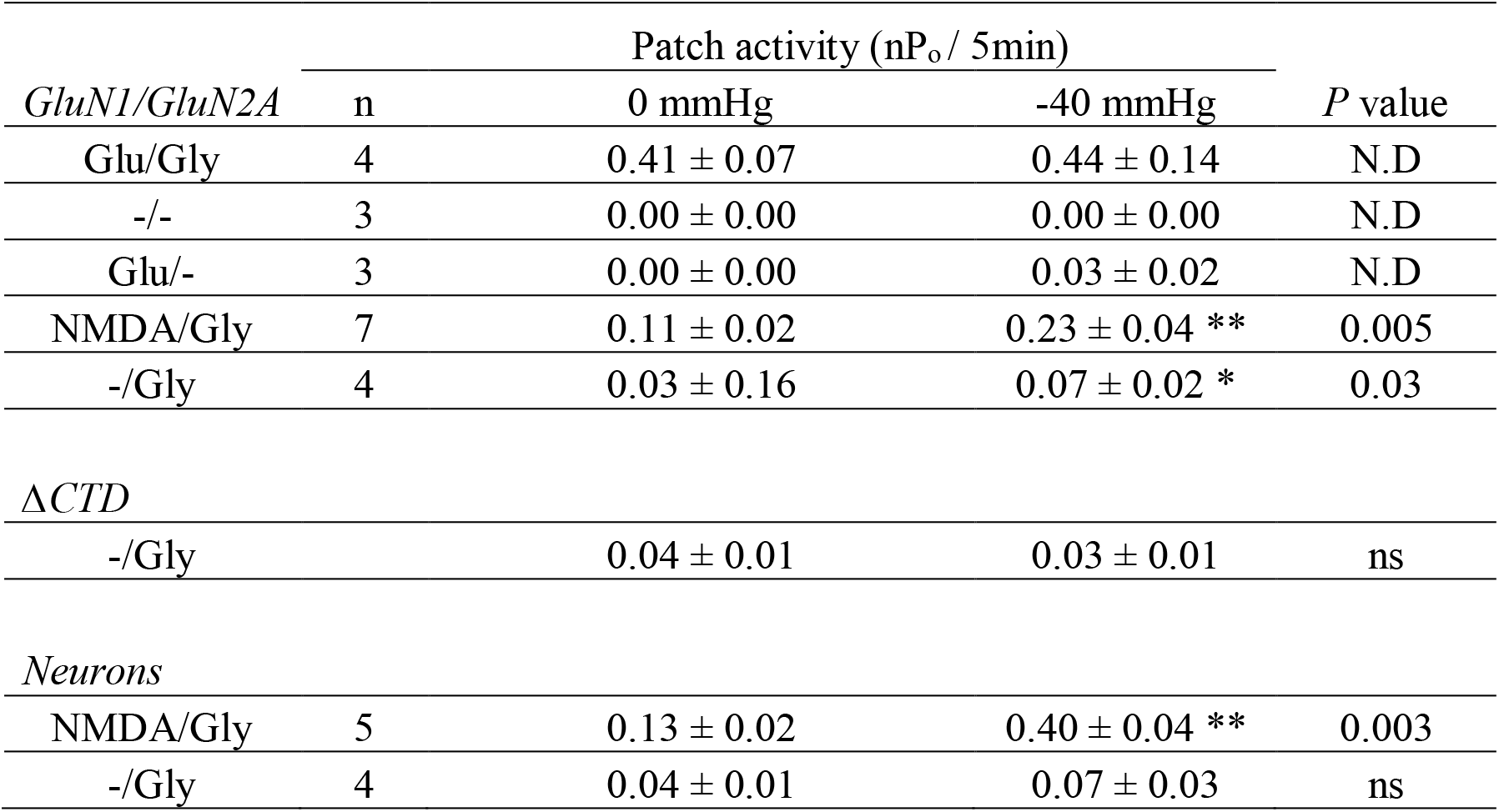
Summary of single-channel activity

|  |  | Patch activity (nPo/ 5min) |  | *P* value |
| --- | --- | --- | --- | --- |
| *GluN1/GluN2A* | n | 0 mmHg | -40 mmHg |  |
| Glu/Gly | 4 | 0.41 ± 0.07 | 0.44 ± 0.14 | N.D |
| -/- | 3 | 0.00 ± 0.00 | 0.00 ± 0.00 | N.D |
| Glu/- | 3 | 0.00 ± 0.00 | 0.03 ± 0.02 | N.D |
| NMDA/Gly | 7 | 0.11 ± 0.02 | 0.23 ± 0.04 ** | 0.005 |
| -/Gly | 4 | 0.03 ± 0.16 | 0.07 ± 0.02 * | 0.03 |
| *ΔCTD* |  |  |  |  |
| -/Gly |  | 0.04 ± 0.01 | 0.03 ± 0.01 | ns |
| *Neurons* |  |  |  |  |
| NMDA/Gly | 5 | 0.13 ± 0.02 | 0.40 ± 0.04 ** | 0.003 |
| -/Gly | 4 | 0.04 ± 0.01 | 0.07 ± 0.03 | ns |

### Biophysical properties of stretch-gated NMDA receptor currents

To estimate conductance and permeability properties of the stretch-gated receptors, after gentle seal formation, we applied −40 mmHg of pressure and recorded activity at several applied pipette potentials for one-minute periods (Figure 3A). From these data, we measured unitary current amplitude at each voltage, and estimated the unitary conductance as the slope of the voltage-current relationship (Figure 3B). Relative to the glutamate-gated currents, which had γ_Na_ = 67 ± 3 pS, stretch-gated currents had a slight but significantly larger unitary conductance, γ_Na_ = 85 ± 7 pS (p < 0.05, unpaired Student’s *t*-test) (Figure 3B). Therefore, stretch-gated currents retain the high unitary conductance characteristic of NMDA receptors.

**Figure 3.**
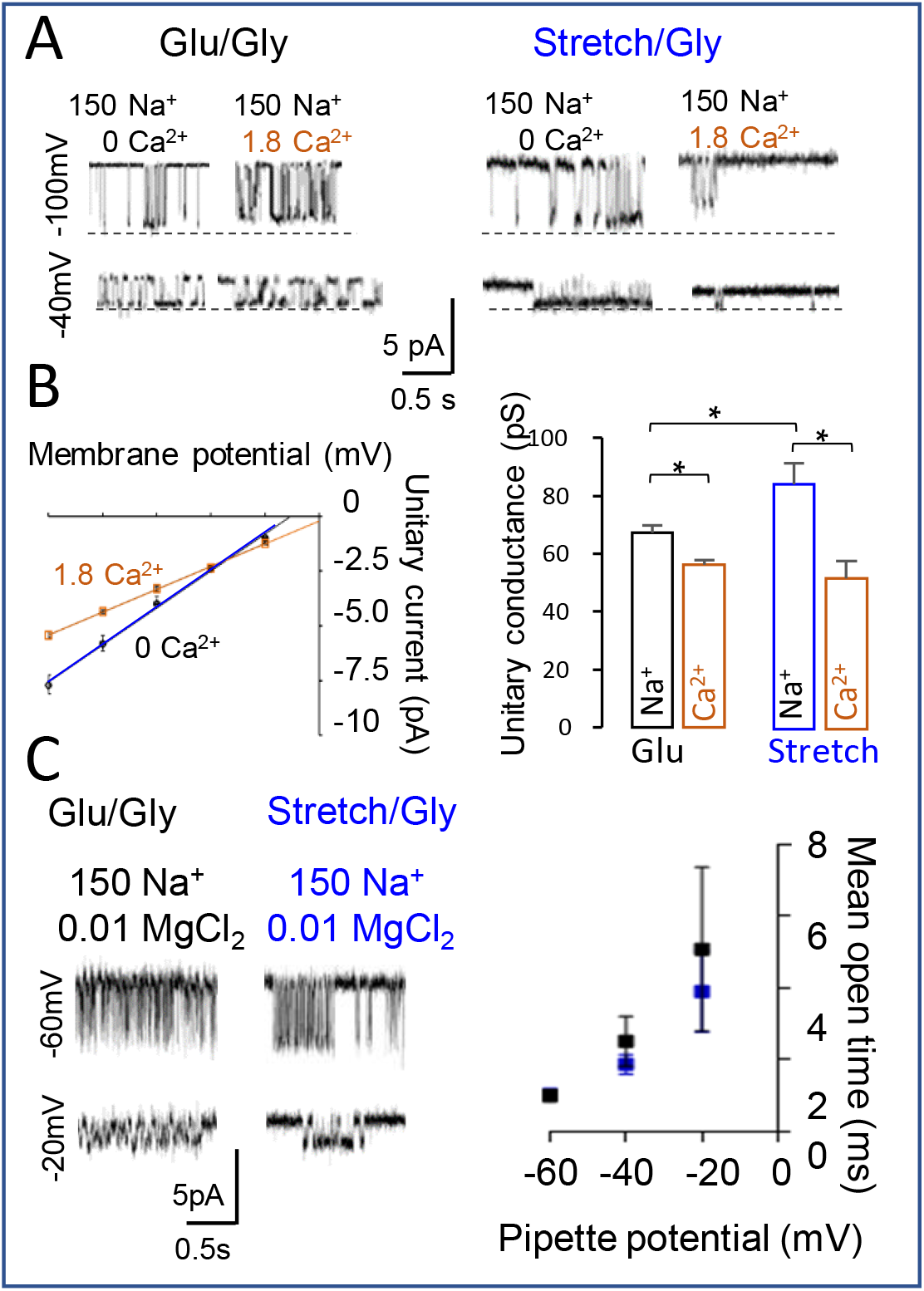
Biophysical properties of stretch-gated currents from recombinant NMDA receptors. **(A)** On-cell patch-clamp current traces recoded from cells expressing GluN1/GluN2A receptors in response to saturating concentrations of Glu (1 mM) (left) and gentle stretch (−40 mmHg). **(B)** Voltage dependency of unitary current amplitude and summary of Ca^2+^-dependent reduction in unitary conductance. **(C)** Current traces recorded with external Mg^2+^ (10 mM) and summary of voltage-dependent reduction in mean open durations.

In physiological conditions, external Ca^2+^ permeates NMDA receptors and also reduces channel conductance. To examine how external calcium affects stretch-gated currents, we measured single-channel current amplitudes of glutamate-gated and stretch-gated currents at several applied voltages, in the absence and presence of external calcium (1.8 mM). We found that 1.8 mM Ca^2+^ reduced the glutamate-gated conductance to γ = 56 ± 2 pS, indicative of ∼25% current blockade (Figure 3B), a value consistent with previous reports (Ascher & Nowak, 1988; B. A. Maki & G. K. Popescu, 2014). Similarly, 1.8 mM Ca^2+^ reduced the amplitude of stretch-gated currents to γ_1.8_ = 52 ± 6 pS. This value is not significantly different from that of glutamate-evoked currents in the same conditions, however it represents a stronger block (∼40%) relative to that observed for glutamate-evoked currents (Figure 3A, B). Therefore, stretch-gated currents remain sensitive to blockage by external calcium, and their unitary conductance in 1.8 mM Ca^2+^ is indistinguishable from that of the glutamate-gated current.

From the same data, we constructed linear fits to the current-voltage relationships obtained in zero and 1.8 mM Ca^2+^, to estimate reversal potentials for each condition. Relative to 0 Ca^2+^, in 1.8 mM Ca^2+^, the reversal potential of glutamate-gated currents shifted by +6 mV, indicative of a high relative Ca^2+^ permeability, P_Ca_/P_Na_ = 6.5, as reported previously (Bruce A. Maki & Gabriela K. Popescu, 2014; Wollmuth & Sakmann, 1998). For stretch-activated currents we measured a +16 mV shift in reversal potential, corresponding to 3-fold increase in permeability P_Ca_/P_Na_ = 18, relative to glutamate-gated currents. Together, these measurements indicate that in physiologic calcium Ca^2+^, stretch gates NMDA receptor currents that maintain characteristic high unitary conductance, voltage-independent Ca^2+^-block, and may have slightly increased Ca^2+^ permeability, when compared with the glutamate-gated currents.

Last, we examined the sensitivity of the stretch-gated current to block by external Mg^2+^. We recorded on-cell single-channel currents from GluN1/GluN2A receptors at several applied voltages, with pipettes containing glycine (0.1 mM), Mg^2+^ (10 µM, Kd = 1 µM) (Premkumar & Auerbach, 1996), and either glutamate (1 mM) or sustained negative pressure (−40 mmHg) (Figure 3C). At each voltage, we identified non-overlapping bursts of activity and measured the channel mean open time as a measure of Mg^2+^-block. As reported previously, we found that glutamate-gated currents were sensitive to block by external Mg^2+^ in a voltage-dependent manner, such that the mean duration of openings decreased from 5.1 ± 2 ms at −20 mV, to 1.0 ± 0.2 ms at −60 mV (Nowak, Bregestovski, Ascher, Herbet, & Prochiantz, 1984). For stretch-gated currents, we observed a similar shortening of open duration with hyperpolarization, from 4 ± 1 ms at −20 mV, to 1.0 ± 0.05 ms at −60 mV (p < 0.05, paired Student’s *t*-test), indicating similar sensitivity to voltage-dependent block (Figure 3C) (Premkumar & Auerbach, 1996). At all examined voltages, the differences in the estimated mean open durations for glutamate-gated and stretch-gated currents were not statistically significant (p > 0.05, unpaired Student’s *t*-test). Together, these results indicate that stretch-activated currents maintain the characteristic biophysical properties of glutamate-gated channels, including high conductance, large Ca^2+^ permeability, strong voltage-dependent Mg^2+^ block, and long openings.

### Stretch-gated NMDA currents require the receptor’s carboxyl terminal

Given the potentially significant physiological implications of a Ca^2+^-rich current gated by mechanical forces through NMDA receptors, it will be important to understand the mechanism by which these arise, and more specifically, to identify the receptor domains responsible for mechanotransduction. The existing literature on the mechanosensitivity of NMDA receptors suggests several mechanisms by which mechanical forces may facilitate the glutamate-gated current. These include a reduction of Mg^2+^ block (Kloda et al., 2007; Mor & Grossman, 2010; L. Zhang, Rzigalinski, Ellis, & Satin, 1996), perhaps transmitted through the transmembrane domain (Casado & Ascher, 1998), but also allosteric mechanisms that implicate the C-terminal domain (Singh et al., 2012). For the stretch-gated current, our results exclude a mechanism mediated by changes in Mg^2+^-block. Therefore, we asked whether the C-terminal domain (CTD) influences the receptor’s mechanically-elicited current.

We recorded single-channel currents from on-cell patches expressing receptors with truncated intracellular C-termini (GluN1^Δ838^/GluN2A^Δ844^). We reported previously that relative to wild-type receptors (WT, Po = 0.54 ± 0.04), glutamate (1 mM) gates currents with substantially lower probabilities from these truncated receptors (ΔCTD, 0.08 ± 0.02, n = 8, P < 0.5) (Maki, Aman, Amico-Ruvio, Kussius, & Popescu, 2012). Using the pressure protocol described here, with only glycine in the pipette and no external pressure, we observed similarly low spontaneous activity from CTD receptors, as reported above for WT (Table 1). However, suction up to −40 mmHg did not increase the basal activity of truncated receptors (Figure 4, Table 1). This result suggests that the CTD of GluN1/GluN2A receptors is necessary for their mechanical activation by gentle suction. This observation may indicate that the CTD is necessary to transmit force from the cytoskeleton to the gate; alternatively, it may indicate that the energy provided by suction, transmitted by some other unknown mechanism, is sufficient to gate the channel only when the tethering of the CTD to intracellular structures provides a certain threshold of rigidity to the observed receptor, a case demonstrated recently for GluD receptors (Carrillo, Gonzalez, Berka, & Jayaraman, 2021).

**Figure 4.**
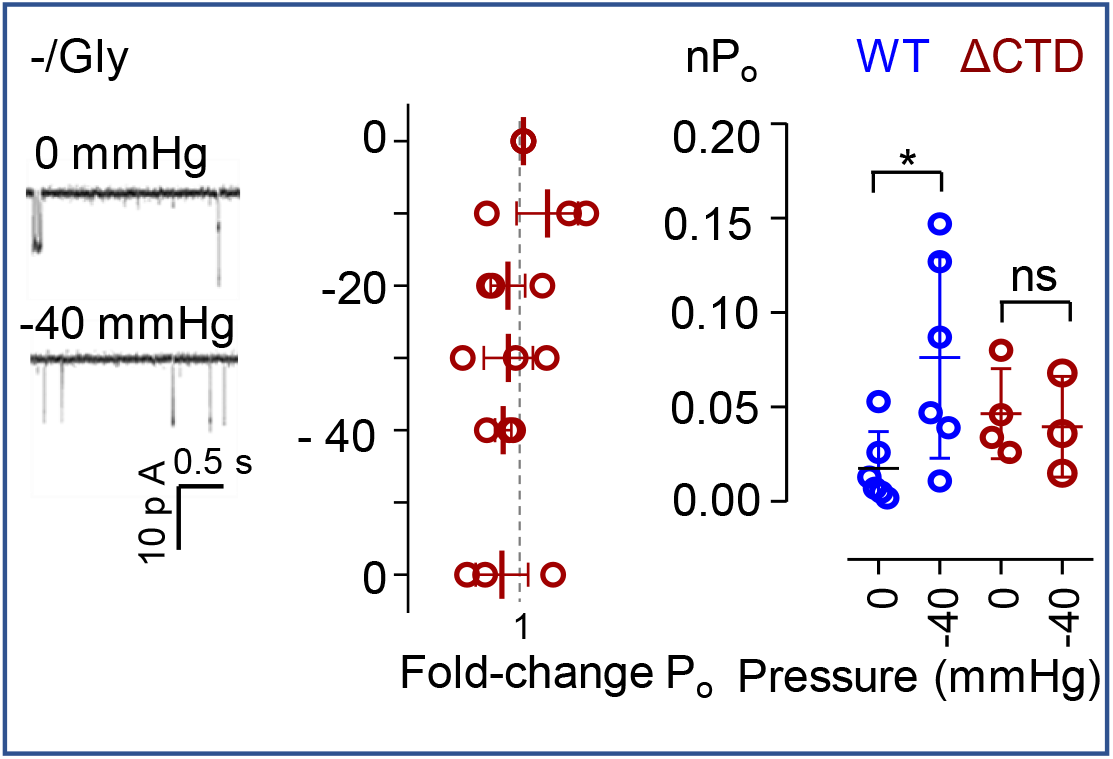
Mechanical activation of NMDA receptors by suction requires their intracellular C-terminal domain. *Left*, Cell-attached Na^+^-current traces recorded from GluN1/GluN2A receptors lacking the intracellular C-terminal domain (ΔCTD). *Right,* summary of results for ΔCTD compared with wild-type (WT) receptors. *, P < 0.05 (paired Student’s t-test).

### Mechanical activation of neuronal NMDA receptors

Regardless of mechanism, our result that CTD is required for mechanical gating of currents from NMDA receptors suggests that the intracellular milieu in which NMDA receptors operate, and specifically the intracellular interactions mediated by the CTD will influence the effectiveness of hydrostatic pressure to gate currents from glycine-bound receptors. In addition, lipid composition of membranes varies widely across cell type, development stage, and subcellular location, and can be a critical determinant of mechanotransduction (Perozo, Kloda, Cortes, & Martinac, 2002; Phillips, Ursell, Wiggins, & Sens, 2009). We therefore investigated the effectiveness of hydrostatic pressure to gate NMDA receptors in a neuronal environment.

We cultured primary rat hippocampal neurons (P7 - P30), and recorded cell-attached currents with pipette solutions containing low concentrations of NMDA (0.1 mM; EC50, 90 mM) (Erreger et al., 2007) and glycine (0.1 mM) to identify currents mediated by endogenous NMDA receptors. We observed inward currents with large unitary amplitudes (8.90 pA ± 0.34) consistent with NMDA receptor activation. Hydrostatic pressure (−40 mmHg) increased substantially the measured nPo from 0.13 ± 0.03 at rest to 0.40 ± 0.05 (n = 8, P < 0.02, Anova post hoc), and this potentiation was fully reversible and mirrored results obtained with low NMDA and glycine fromGluN1/GluN2A receptors in HEK293 cells (Figure 5A, Table 1). In similar experiments and with only glycine in the pipette, hydrostatic pressure (−40 mmHg) increased the observed nPo from 0.04 ± 0.01 at rest, to 0.07 ± 0.02, (n = 5, P < 0.05, Anova post hoc test), an activation which was reversible upon returning to resting conditions (Figure 5B, Table 1).

**Figure 5.**
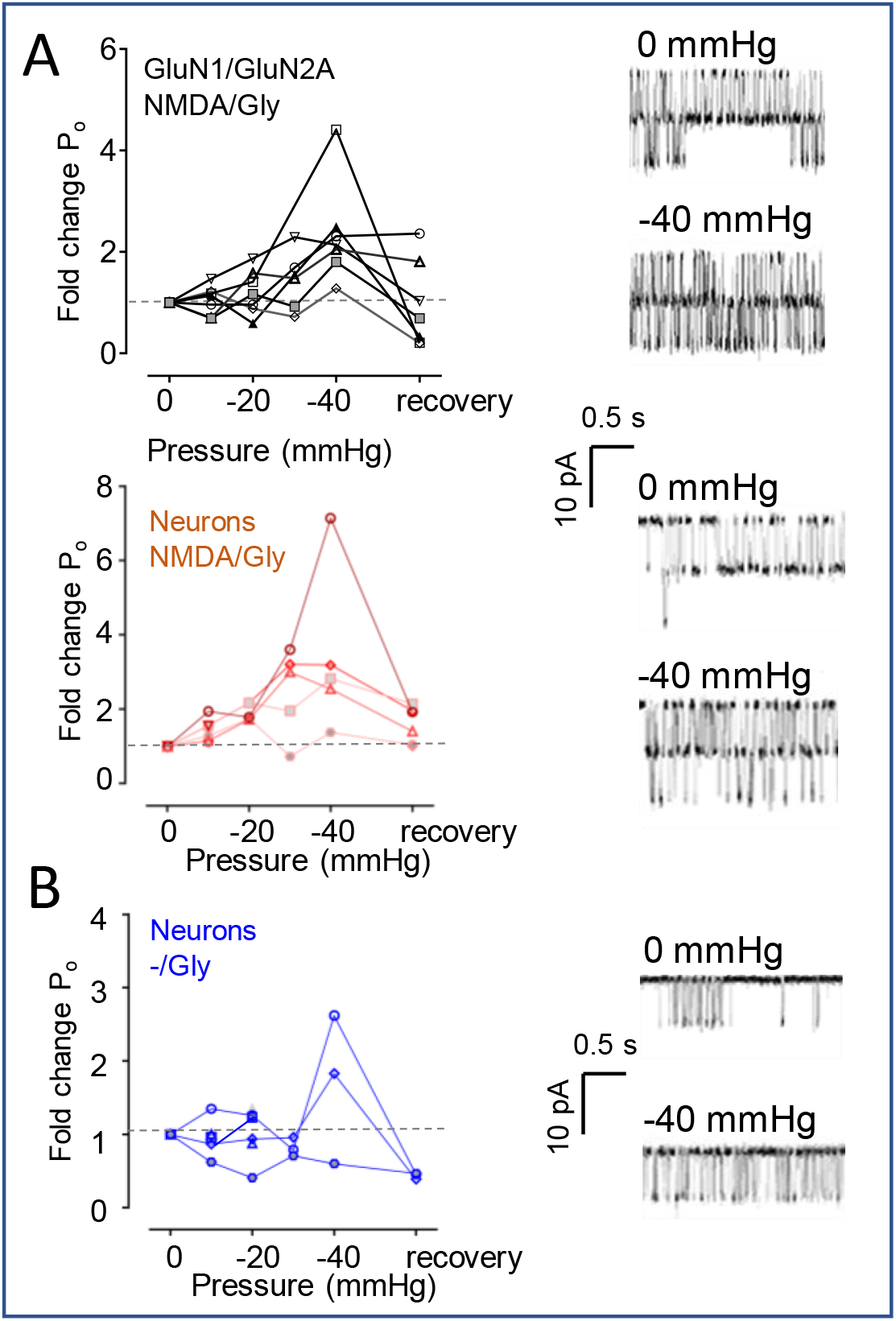
Gentle membrane stretch gates native NMDA receptors. **(A).** Suction potentiates currents elicited with low concentration of NMDA (0.1 mM) from recombinant GluN1/GluN2A receptors (HEK cells) and from native NMDA receptors in cultured rat hippocampal neurons. **(B).** Gentle suction gates NMDA receptor-like currents from cultured hippocampal neurons.

Together, these observations validate the results obtained with recombinant receptors in HEK cells and support our proposal that mild stretch can gate native NMDA receptors in the absence of neurotransmission, and likely potentiate responses elicited by low concentrations of glutamate (< 0.1 mM), as may occur at extrasynaptic locations (Moldavski et al., 2020).

## Discussion

Glutamate-gated NMDA receptor currents can be modulated by several types of mechanical perturbations including those generated by changes in environmental pressure (Fagni et al., 1987; Mor & Grossman, 2006), membrane composition (Casado & Ascher, 1998; Miller et al., 1992; Nishikawa et al., 1994), osmotic and hydrostatic pressure (LaPlaca & Thibault, 1998; Paoletti & Ascher, 1994), and microfluidic sheer stress (Maneshi et al., 2017). Although the effects of mechanical stimulation on channel responses vary across stimulation procedure, receptor preparations, and experimental conditions, these results have established that NMDA receptors are mechanosensitive (Paoletti & Ascher, 1994). Here, we report that gentle suction can activate NMDA receptors in the absence of glutamate, such that 40 mmHg of negative pressure produces ∼15% of the maximal glutamate-gated response. This observation establishes that NMDA receptors, in addition to being mechanosensitive, can be activated mechanically, which is consequential to understanding the mechanobiology of the central nervous system. Before addressing this point of impact, we note a number of caveats.

As previously reported for NMDA receptor mechano-sensitivity (Paoletti & Ascher, 1994), the mechano-activity we observed here for NMDA receptors was variable, despite taking a number of experimental precautions. Among these, to ascertain whether NMDA receptors can be activated by mechanical forces, we examined the effect of applied hydrostatic pressure onto recombinant receptors residing in cell-attached membrane patches. This approach has a number of advantages. The cell-attached configuration maintains cellular integrity and a near-physiologic cellular environment for the examined receptor; it allows precise control of the magnitude of the applied pressure with a high-speed pressure clamp; and provides a high-resolution single-molecule readout for receptor activity. Despite these advantages, a direct correlation between the applied pressure and the receptor’s microscopic properties is complicated by a number of uncontrollable variables. First, even when using pipettes of specified geometry and resistance, the area of the electrically accessible membrane patch (the dome delimited by the seal) varies from patch to patch and can change during a single recording due to membrane creep (Suchyna, Markin, & Sachs, 2009). Further, the tension experienced by receptors varies with their position within the patch, being highest at the apex of the patch and lowest near the pipette wall (Bavi et al., 2014). Lastly, the size and mechanical properties of the cytosolic plug that accompanies the patch within the pipette is not uniform across experimental observations, and furthermore it can detach from the bilayer upon continuous mechanical stimulation. Such blebbing may produce additional variability in the magnitude of the force that reaches the receptor and can also modify the receptor’s gating properties (Krupp, Vissel, Thomas, Heinemann, & Westbrook, 1999a; Suchyna et al., 2009). Although these sources of variability apply to the study of other channels, for which combining pressure-clamp and cell-attached electrophysiology was successful, they may be particularly obtrusive when examining NMDA receptors, which have notoriously complex reaction mechanisms (Iacobucci & Popescu, 2017).

At the single-channel level, NMDA receptors present intrinsic gating heterogeneity due to modal gating (G. Popescu & Auerbach, 2003; G. K. Popescu, 2012; Vance et al., 2013), which results in a characteristic biphasic decay of their macroscopic response (W. Zhang, Howe, & Popescu, 2008). For this reason, even in controlled experimental conditions, the measured equilibrium open probability of NMDA receptors varies considerably (Borschel et al., 2012; Vance et al., 2013). In addition, NMDA receptor activity is sensitive to cellular factors that may vary from cell to cell and may change during extended recording periods (Cerne, Rusin, & Randic, 1993; Chen & Huang, 1991; Tong, Shepherd, & Jahr, 1995; Wyszynski et al., 1997). With these considerations in mind, the magnitude of the changes in activity we observed with gentle suction are consistent with substantial mechano-activation of the four NMDA receptor isoforms examined here.

This observation is important for several reasons. Regulated NMDA receptor-mediated calcium influx is required for the normal development and function of excitatory synapses, and mechanical forces may be important in initiating these processes during development and throughout the life span (Tyler, 2012). Alternatively, NMDA receptor-mediated calcium can also initiate synaptic pruning, spine shrinkage, and neuronal death. NMDA receptors are expressed not only at post-synaptic, mechanically stable locations, but also in mechanically active or osmotically sensitive zones, such as growing axons or dendritic boutons, where local deformations in extracellular matrix, membrane tension or curvature, and intracellular cytoskeleton can impinge on receptors to change their conformational landscape. Therefore, NMD receptors operate in a mechanically rich landscape and depending on their location may be more or less exposed to changing mechanical forces. Our results show no effect of membrane stretch on currents elicited with maximally effective glutamate concentrations (Figures 1A and 2, and Table 1). Therefore, it is unlikely that this mechanism will influence synaptic transmission. However, the levels of mechano-activation we observed with gentle membrane stretch can have a significant impact on signal transduction by neuronal extrasynaptic NMDA receptors, or those expressed in glial cells. Additionally, NMDA receptors, of unknown function, have been identified at non-traditional sites such gastrointestinal, lung, and adrenal tissue during human development (Szabo et al., 2015); and adult sites such as kidney (Leung et al., 2002), bone (Itzstein et al., 2001), myocytes (Dong et al., 2021; Seeber et al., 2004), colon (Del Valle-Pinero et al., 2007) and others, such as cancerous tissue (Yan et al., 2021). Therefore, the significance of the mechano-activity described here will vary with the site of NMDA receptor expression and their microenvironment.

For the most prevalent NMDA receptor isoform expressed in adult mammals, GluN2A-containing receptors, −40 mM Hg produced 15% of the maximal glutamate-elicited activity. In pharmacology, this observation would indicate that hydrostatic pressure is a partial agonist at the glutamate site of the NMDA receptors. Because we did not test pressures above 40 mmHg, and because when using the more sensitive protocol, the response did not appear to plateau (Figure 2), it is possible that stronger forces may elicit higher activity. However, we can expect that the 15% activation of NMDA receptors we observed with this gentle level of membrane stretch, which can be assumed to occur endogenously during physiologic conditions, represents a biologically significant level of signal transduction. The ambient glutamate concentration at extrasynaptic sites in adult rat hippocampal slices is estimated at 25 - 80 nM (Herman & Jahr, 2007; Moldavski et al., 2020), which represents less than 10% of the synaptic concentration (∼ 1 mM) (Budisantoso et al., 2012; Clements, Lester, Tong, Jahr, & Westbrook, 1992; Wadiche & Jahr, 2001). Together with the observation reported here and previously (Casado & Ascher, 1998; Paoletti & Ascher, 1994) that gentle membrane stretch potentiates responses elicited with low concentrations of the GluN2-site agonist (glutamate or NMDA), the mechano-activity of NMDA receptors may represent an important physiologic mechanism, especially in development or at sites of dendritic growth and synaptic formation. Alternatively, inappropriate mechanical activation of extrasynaptic NMDA receptors, due to for example external mechanical forces experienced by brain or spinal cord, may initiate or aggravate apoptotic or necrotic cell injury through additional Ca^2+^ influx.

The precision with which we measured the mechanical activation of GluN2B-, GluN2C- and GluN2D-containing receptors was insufficient to estimate the efficacy of stretch relative to glutamate for these isoforms. Moreover, even with glutamate as the agonist, these receptors have much lower maximal open probabilities: 0.16 ± 0.02 for GluN2B (Borschel et al., 2012); 0.032 ± 0.015 for GluN2C (Khatri et al., 2014), and 0.023 ± 0.001, for GluN2D (Vance et al., 2013), and 0.07 to 0.39 for neuronal receptors (Borschel et al., 2012). Nonetheless, our results show that mechanical activation of NMDA receptors can produce substantial mechanotransduction, even when glutamate is absent. Together with previous reports that mechanical forces facilitate the current gated by low concentrations of glutamate, indicate that NMDA receptors represent important mechanotransducers in the CNS. However, the mechanism by which NMDA receptor sense mechanical forces and how these are transmitted to the channel gate remains to be determined.

In some experimental paradigms the mechanosensitivity of NMDA receptors reflects mechanically-induced changes in the receptor’s sensitivity to voltage-dependent Mg^2+^ block (Cox et al., 2019; Kloda et al., 2007; Parnas et al., 2009; L. Zhang et al., 1996). Given that the majority of our measurements were done in the absence of external Mg^2+^, and we were able to demonstrate similar voltage-dependent block for stretch-gated and glutamate-gated currents (Figure 3C), we can definitively exclude this mechanism for the stretch-gated activity we examined here. Aside from modulating Mg^2+^-block, previous reports found mechanosensitivity to depend on the receptor’s intracellular CTD (Bliznyuk, Aviner, Golan, Hollmann, & Grossman, 2015; Singh et al., 2012). Our results show that the CTD of NMDA receptors is required for mechanical activation by gentle membrane stretch.

Given the modular make-up of NMDA receptors, and their complex interaction with extracellular matrix proteins, membrane proteins and lipids, and with intracellular proteins and cytoskeletal components, it is likely that depending on the type of stimulation, mechanical forces will impinge on separate receptor domains. For example in the experiments reported by the Martinac group (Kloda et al., 2007), when mechanosensitivity was tested on purified recombinant NMDA receptors inserted in liposomal particles, it was reasonable to infer a force-from lipid transduction mechanism, given the absence of interacting proteins or cellular structures. However, when operating in their native environments, receptors are much more mechanically constrained and they can sense membrane deformation not only through direct interactions with lipid but also through their extracellular or intracellular domains. In addition, mechanical constraints imposed by interaction with cellular and/or intracellular structures, even if not serving as mechanical transducers, may serve to limit the receptor’s conformational freedom and thus facilitate or oppose their mechanical activation. This scenario was recently described for GluD receptors, which can be gated by neurotransmitter only when their N-terminal domains are reinforced by interactions with trans-cellular proteins (Carrillo et al., 2021). Therefore, although the CTD appears necessary for mechanical gating of NMDA receptors, it remains to be determined whether this is a case of force-from filament mechanism, or the CTD simply stabilizes the molecule in a mechano-sensitive conformation.

Therefore, although unknown at this time, it will be important to determine the mechanism by which mechanical forces gate NMDA receptors currents. Mechano-activation of NMDA receptors may be of particular importance at extrasynaptic and non-neuronal sites in the CNS, where it may contribute to fundamental processes such as synapse formation, dendrite remodeling, and glial physiology, and outside of the nervous system in gastric and pulmonary development, and cardiac and bone remodeling. Conversely, this knowledge may help better understand, and therefore prevent or address, neuropsychiatric syndromes such as shaken baby, chronic traumatic encephalopathy, and other trauma-associated neuropathologies.

## Acknowledgements

This work was supported by R21NS098385, R01NS052669 and R01NS097016 to GKP. We thank Eileen Kasperek for assistance with molecular biology and tissue culture. We thank Richard Burke, Cheryl Movsesian, and Ayman Mustafa for sharing recordings. We thank Gary J. Iacobucci for his assistance in developing the Matlab code used to analyze the SKM data.

## Author contributions

S.B., B.A.M., and G.K.P. contributed to study design and figure preparation; S.B., B.M., J.C., and B.R acquired and analyzed data, S.B. and G.K.P. wrote the article.

## Notes

### Competing Interest Statement

The authors have declared no competing interest.

